# Functional MRI reveals memories of mother in newborn chicks

**DOI:** 10.1101/2022.02.12.480179

**Authors:** Mehdi Behroozi, Elena Lorenzi, Sepideh Tabrik, Martin Tegenthoff, Alessandro Gozzi, Onur Güntürkün, Giorgio Vallortigara

## Abstract

Filial imprinting has been used as a powerful ethological paradigm to investigate the neurobiology of early learning that affects lifelong behaviours. When a visually naïve chick is exposed to one of a wide range of conspicuous objects, it may learn its characteristics and subsequently recognises and selectively approaches this stimulus (usually the mother hen and siblings). While the initial phases of memory acquisition have been unravelled, the long-term storage and retrieval components of imprinting memories are still unknown. Here, we used functional MRI in awake newly hatched chicks to identify the long-term storage of imprinting memories in a wide network comprising the hippocampal formation, the medial striatum, the arcopallium, and the prefrontal-like nidopallium caudolaterale. These results provide the first example of non-invasive imaging of the brain in a newborn vertebrate.

## Introduction

Filial imprinting is a learning process by which the young of some organisms can learn about a conspicuous object, usually the mother or siblings, by simply being exposed to it for a short period of time soon after birth (McCabe, 2013). It owes its great popularity to the work of Nobel-prize-winning ethologist Konrad Lorenz (Lorenz, 1935), but it was originally described by Douglas Spalding (Spalding, 1954) in the offspring of some nidifugous (precocial) bird species, such as chicks or ducklings (see Vallortigara, 2021). Visual imprinting has been mostly studied, though acoustic or olfactory imprinting can be observed as well, the latter being prominent in mammals (Müller-Schwarze and Müller-Schwarze, 1971).

Although in principle visual imprinting can occur with any kind of object, research has shown that the process is actually assisted by a set of biological predispositions which guide an animal’s attention towards those object features that are more likely to be observed in social partners - e.g., preferences in domestic chicks include simple features such red colour (which is prominently observed in the head region of conspecifics), or self-propelled motion (which is typical of living things), as well as more complex assembly of features such as face-like stimuli or biological motion in point-light displays (review in Lorenzi and Vallortigara, 2021; Vallortigara, 2021). Brain research has shown that biological predispositions are associated with activation of areas of the so-called *Social Behavior Network*, and in particular of the lateral septum for motion stimuli and of the nucleus taenia (homologous of the mammalian medial amygdala (Yamamoto et al., 2005)) for face-like stimuli (review in Lorenzi and Vallortigara, 2021).

Interest for filial imprinting quickly spanned from behavioural biology to psychological development and psychopathology, inspiring for instance John Bowlby’s theory of attachment, which postulates a crucial role of the mother-child bond for subsequent psychological development and, complementarily, the psychiatric outcomes associated with early mother deprivation (recent reviews in Lemche, 2020; Vicedo, 2013).

In the 70’s filial imprinting served as a model for the neurobiological investigation of memory. Gabriel Horn and colleagues (review in Horn, 2004) identified an associative brain region involved in the formation of an imprinting memory, the intermediate medial mesopallium (IMM according to the new avian brain nomenclature; previously referred to as IMHV, intermediate medial hyperstriatum ventrale (Jarvis et al., 2005; Reiner et al., 2004)). IMM proved to be crucial during the acquisition phase of the visual imprinting memory. More precisely, it was found that exposure to the imprinting object was associated with changes in the left but not in the right IMM (Bradley et al., 1981; Horn et al., 1985). Subsequent studies with auditory imprinting revealed that the imprinting-related area extended ventrally into a medialmost nidopallial area, the nidopallium medial pars medialis (NMm) (Bredenkötter and Braun, 2000, 1997). Here we will use the label medial nidopallium/mesopallium (MNM) to jointly label the mesopallial and nidopallial entity of the imprinting area.

Experiments involving sequential lesions, first to one side of IMM and subsequently to the other (Cipolla-Neto et al., 1982; Horn et al., 1983), suggested that the store in the left IMM is only temporary, and the right IMM is implicated in transferring information from the left IMM to another, unknown brain region dubbed S’, and that this transfer appears to be complete within 6 h after the end of exposure (Davey et al., 1987). Thus, to cite Gabriel Horn’s words *“We are still some way from being able to visualize, through the microscope or by using brain imaging techniques, the neural trace of (imprinting) memory”* (Horn, 2004). Here we set out to find an answer to this open question.

Techniques such as functional magnetic resonance imaging (fMRI) would allow for non-invasive investigation of ongoing brain activity and look ideally-suited for the time-course of the formation of an imprinting memory. Here we therefore devised an fMRI study in awake newly-hatched chicks to identify the imprinting network and the long-term store of imprinting memories.

We exposed (imprinted) chicks on either a preferred (red) or a non-preferred (blue) colour (Maekawa et al., 2006; Salzen et al., 1971). After exposure, chicks were tested with a sequence alternating the two colours in the scanner. Chicks imprinted on red colour showed activity in pallial and subpallial brain regions involved with storage and memory retrieval, such as the medial striatum, the arcopallium, the hippocampus, and the nidopallium caudolaterale (a presumed avian equivalent of mammalian prefrontal cortex). Chicks imprinted on blue showed little or no activity in the same regions but, as a result of exposure in the scanner to the preferred red colour, started a process of secondary imprinting, which confirmed an early activation of mesopallium, as well as a precocial involvement of the *Social Behavior Network* during the first exposure to a predisposed feature, such as the colour red. We thus for the first time were able to image both the brain areas involved in the acquisition of the imprinting memory, with associated areas involved in biological predispositions that canalise the learning process, and the areas – unknown up to now - associated with long-term store of imprinting memory.

## Results

### Establishment of a fully non-invasive and awake fMRI for the chicks

To enable whole-brain fMRI acquisition in awake chicks, we developed a fully non-invasive set-up to minimise head and body movements (Fig. 1B). Before fMRI scans, chicks were imprinted for two days on either a preferred red or a non-preferred blue light ball (Salzen et al., 1971). At first, we tried to habituate newborn chicks to the head fixation system before the scanning session. However, the habituation sessions stressed the animals so much that ended up panicking against the head fixation system already at the second habituation session. Therefore, we did not habituate the animals to the head fixation before the scan, in order to avoid any association between stress or anxiety and the head fixation system. So before scanning, chicks were only habituated to the scanner noise (Fig. 1A). On the third day, after wrapping the animal in a paper tissue to avoid any body-part movements (such as wings and legs), blocks of plasteline were used to comfortably fixate the head, minimising movements and scanner’s noise by covering their ears (Fig. 1B).

**Fig.1.**
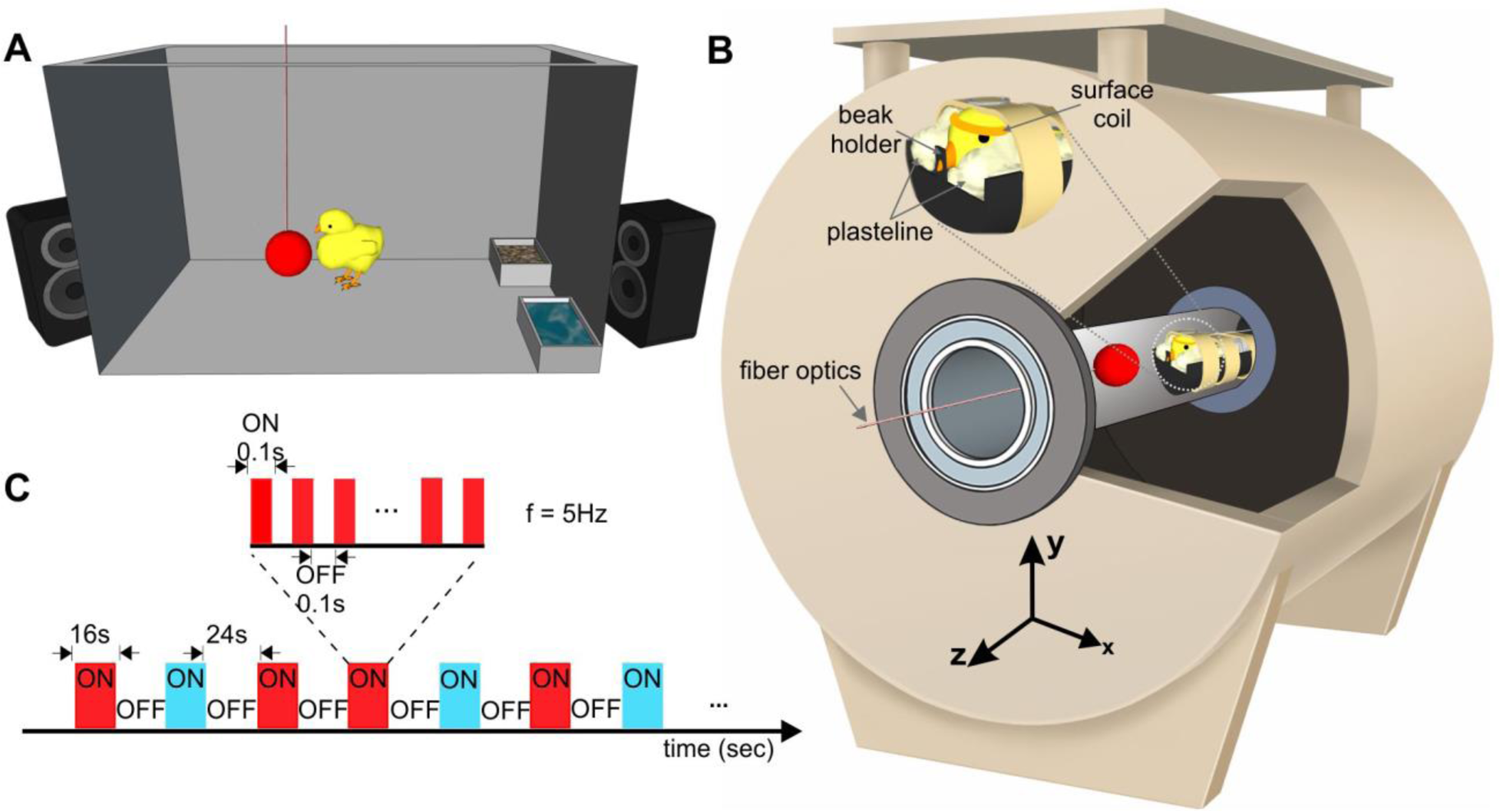
Experimental setups and stimulation sequence for awake chick fMRI. (A) Imprinting cage. Newborn chicks were first exposed to a hollow plastic ball with a flickering red/blue light at a frequency of 5Hz. (B) Custom-made restrainer and 7T fMRI system. Awake chicks were placed in an MR-compatible tube. To immobilise non-invasively the animals, a beak holder was used to control the beak movements, and blocks of plastelines were used to cover the ears and reduce head movements. To avoid body-part movements, animals were wrapped in paper tissue before fixating the head. Subsequently, the animal’s body was taped to the restrainer. (C) A sequence of the block design experiment paradigm. Visual stimuli were presented in blocks of 16 s followed by 24 s dark. During the ON blocks, the visual stimulus (red/blue light) flickered at a frequency of 5Hz.

To record the spontaneous resting-state and task-based BOLD signals, a single-shot multi-slice rapid acquisition with relaxation enhancement (RARE) sequence was adopted from Behroozi et al (Behroozi et al., 2020, 2017). Voxel-wise signal-to-noise ratio (SNR) and temporal SNR (tSNR) were calculated over the resting-state (rs-fMRI) and task-based fMRI (tb-fMRI) scans respectively. The tSNR of the RARE sequence in each voxel was calculated after applying motion correction and high-pass temporal filtering (cut-off at 120s) to remove any linear drift. Temporal SNR in the entire telencephalon ranged from 50 to 100 (Fig. S1A, B) for both tb- and rs-fMRI scans. Furthermore, the result indicated highly correlated SNR and tSNR for both rs- and tb-fMRI scans (Fig. S1C, D).

In order to verify that adequate fixation was achieved during fMRI scans, we used the realignment parameters and the results of the frame-wise displacement (FD) to evaluate the amount of head motion. Overall, the custom-made restrainer yielded a low level of head movements. There was only 2.02 % (218 volume) and 1,08% (19 volumes) of fMRI volumes with FD higher than 0.2 mm (∼40% of voxel size) over all subjects in the task-based and resting-state experiments, respectively (Figure 2A). The median of frame-wise displacement was ∼0.03 mm for both tb-fMRI and rs-fMRI experiments. However, most head movements occurred in the y-direction (Figure 2B, C). The respective violin plot information for translations in y-direction is as follow: tb-fMRI: max/min = 0.22/-0.31 and median ∼ 0; and rs-fMRI: max/min = 0.28/-0.30 and median ∼ 0. The higher motion parameters in the y-direction were most likely due to the design of the head restrainer, which allowed movements in the dorsoventral direction to avoid blocking the throat.

**Fig.2.**
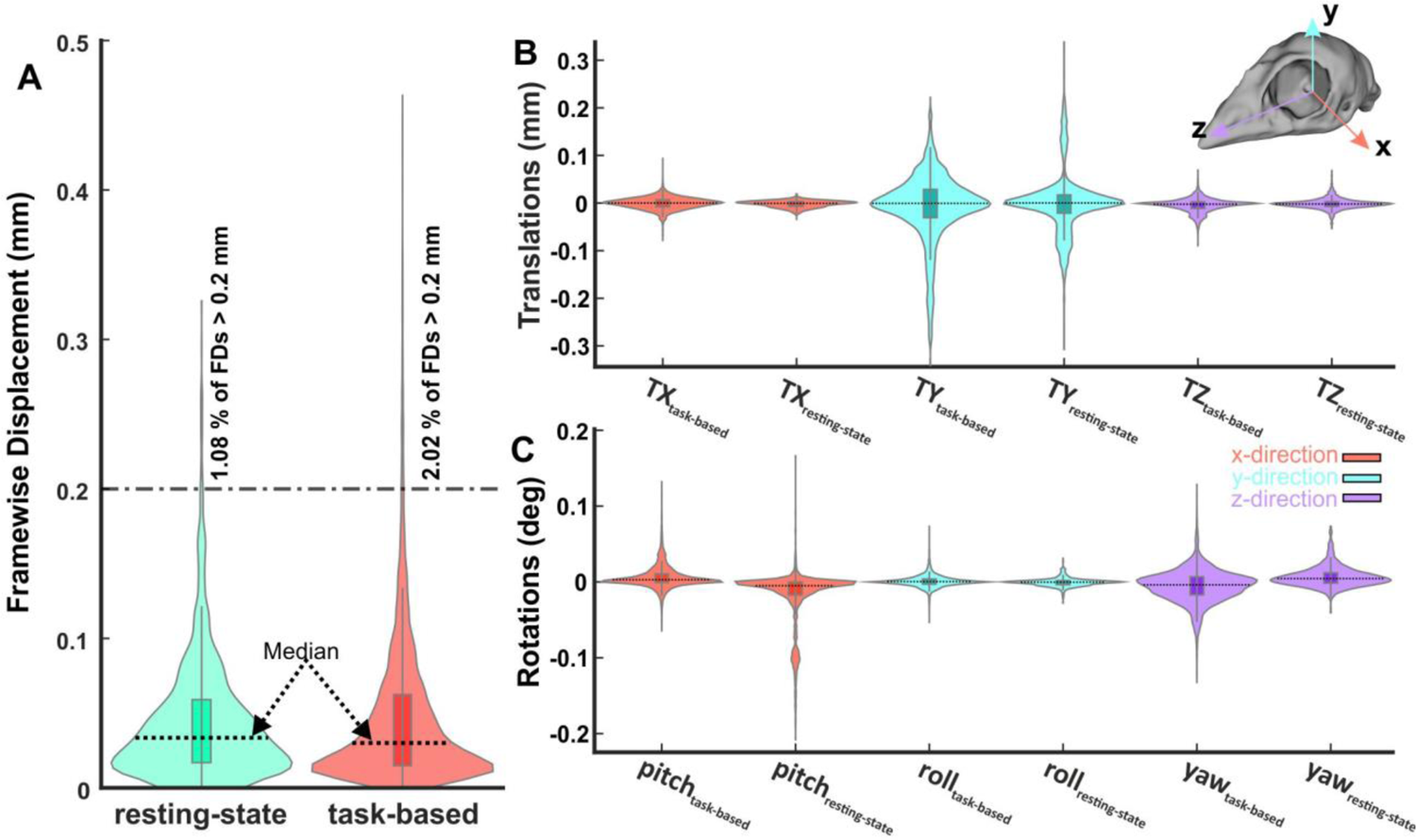
Characterisation of the head movements. (A) Violin plot of the frame-wise displacement over all subjects for task-based (n = 17) and resting-state (n = 9) fMRI scans, respectively. The median is represented by a dash-line across the kernel density estimate of the data. (B, C) Violin plot of estimated motion parameters. Motion parameters were estimated by a 3D rigid body model with six degrees of freedom for translation (x, y, and z-direction) and rotation (pitch, roll, and yaw). The median is represented by a dash-line across the kernel density estimate of the data.

### Distinct BOLD response to identify the acquisition and long-term storage of imprinting memory

We recorded the whole brain BOLD signals from 17 head-restrained awake chicks already imprinted to either a preferred colour, red (n = 9), or a non-preferred colour (Maekawa et al., 2006; Salzen et al., 1971), blue (n = 8). During fMRI scanning, animals were presented with both colours (Fig. 1C), which depending on the previous imprinting training could represent either the imprinted or the control colour. The two colours were presented in a block design manner and a pseudo-random order (48 trials, 24 per condition, see Methods). For the preferred colour group, the imprinting colour (Imp) was red and the control (Cont) was blue, while for the non-preferred colour group the imprinting colour was blue and the control red. To identify the long-term storage of imprinting memory, we used the contrast of Imp > Cont in a conventional generalised linear model (GLM) based statistical analysis. The first-level results at the single-subject level were then entered into a second-level analysis (random-effect modelling, Z = 3.1 (red group) and p < 0.05 family-wise error (FWE)) to illustrate the activation clusters at different networks of chick prosencephalon.

Before fMRI scans, chicks were exposed to the imprinting stimulus for 2 days, during which they learned the feature of the imprinting object and stored them as a long-term memory (McCabe, 2013). Therefore, we expected to find activation in regions involved in memory retrieval. While Imp > Cont contrast in the red group showed robust activation clusters in many telencephalic as well as diencephalic regions, Imp > Cont contrast in the blue group showed no significant activation clusters.

This is due to chicks’ preference for red over blue (Maekawa et al., 2006), therefore we used the Cont > Imp contrast (red > blue colour) during the last 20 minutes of scanning, to investigate the memory formation phase of a secondary imprinting process (Boakes and Panter, 1985) elicited by the presence of the preferred colour red. As illustrated in Figure 3, the voxel-based group analysis showed robust BOLD responses in different visual prosencephalic regions: the nucleus geniculatus lateralis pars dorsalis (GLd, which receives direct input from the retina (Güntürkün and Karten, 1991)), the right intermediate hyperpallium apicale (IHA, which primarily receives visual thalamic input (Csillag and Montagnese, 2005)), the right hyperpallium intercalatum (HI) and right hyperpallium densocellulare (HD), and bilaterally the hyperpallium apicale (HA, together with HD associative hubs of the thalamofugal pathway (Csillag and Montagnese, 2005; Shanahan et al., 2013)) of the thalamofugal pathway, bilaterally the nucleus rotundus (Rot, which is the primary thalamic input region of the tectofugal pathway). Also parts of the auditory system were activated: bilaterally the ventromedial part of the Field-L complex and the right nucleus ovoidalis (OV), a thalamic auditory nucleus receiving direct input from the avian homologue of the inferior colliculus (*torus semicircularis* (Karten, 1967)) that projects to Field-L. We detected significant activation clusters in the associative pallial regions nidopallium medial pars medialis (NMm) and bilaterally in the caudal intermediate medial mesopallium (IMM). Within the two interconnected *Social Behavior Network* and *Mesolimbic Reward System*, we detected a significant BOLD increase rightward in the bed nucleus of the stria terminalis (BNST), the nucleus accumbens (Ac) and the medial striatum (MSt), bilaterally in the septum and leftward in the posterior pallial amygdala (PoA) and the ventromedial part of hippocampus (Hp-VM).

**Fig.3.**
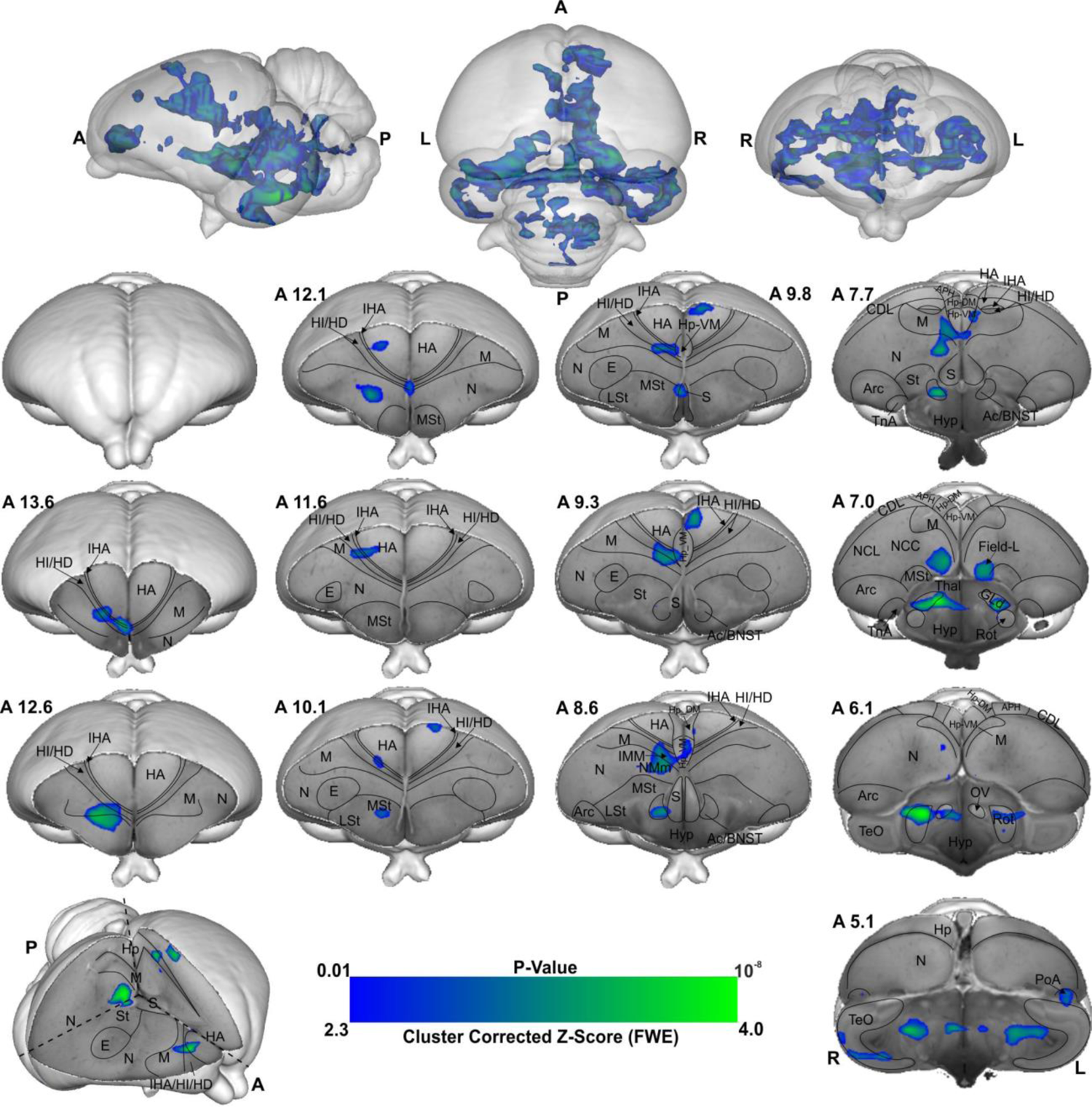
BOLD response pattern during the acquisition of imprinting memory. Statistical maps for the BOLD signal increase in the contrast of red light versus blue light in the Blue group (n = 8, Z = 2.3, and p < 0.05 FEW corrected at the cluster level). The top row images show the 3D representation of the activation pattern inside a translucent chick brain. A 3D depiction of the chick brain is represented at the bottom of the left column with an example window at the level of A 7.0. Anatomical borders (black lines) are based on the contrast difference of ex-vivo chick brain and Chick atlas (Herold et al., 2014; Kuenzel and Masson, 1988). The corresponding abbreviations of ROIs are listed in the Table S1.

As illustrated in Figure 4, the voxel-based group analysis during the imprinting memory retrieval phase in the red group showed robust BOLD responses in different visual prosencephalic regions: the right GLd, bilaterally in IHA, HI, HD and HA (Fig. 4). We found also a significant BOLD rightward increase in part of the auditory system, OV. Furthermore, we detected a significant increase in the BOLD signal in the associative right MNM (IMM + NMm) and nidopallium caudolaterale (NCL) and in left portions of the caudal mesopallium dorsale (MD) and nidopallium caudocentrale (NCC; all interconnected regions (Aoki et al., 2015; Atoji and Wild, 2009; von Eugen et al., 2020)).

**Fig.4.**
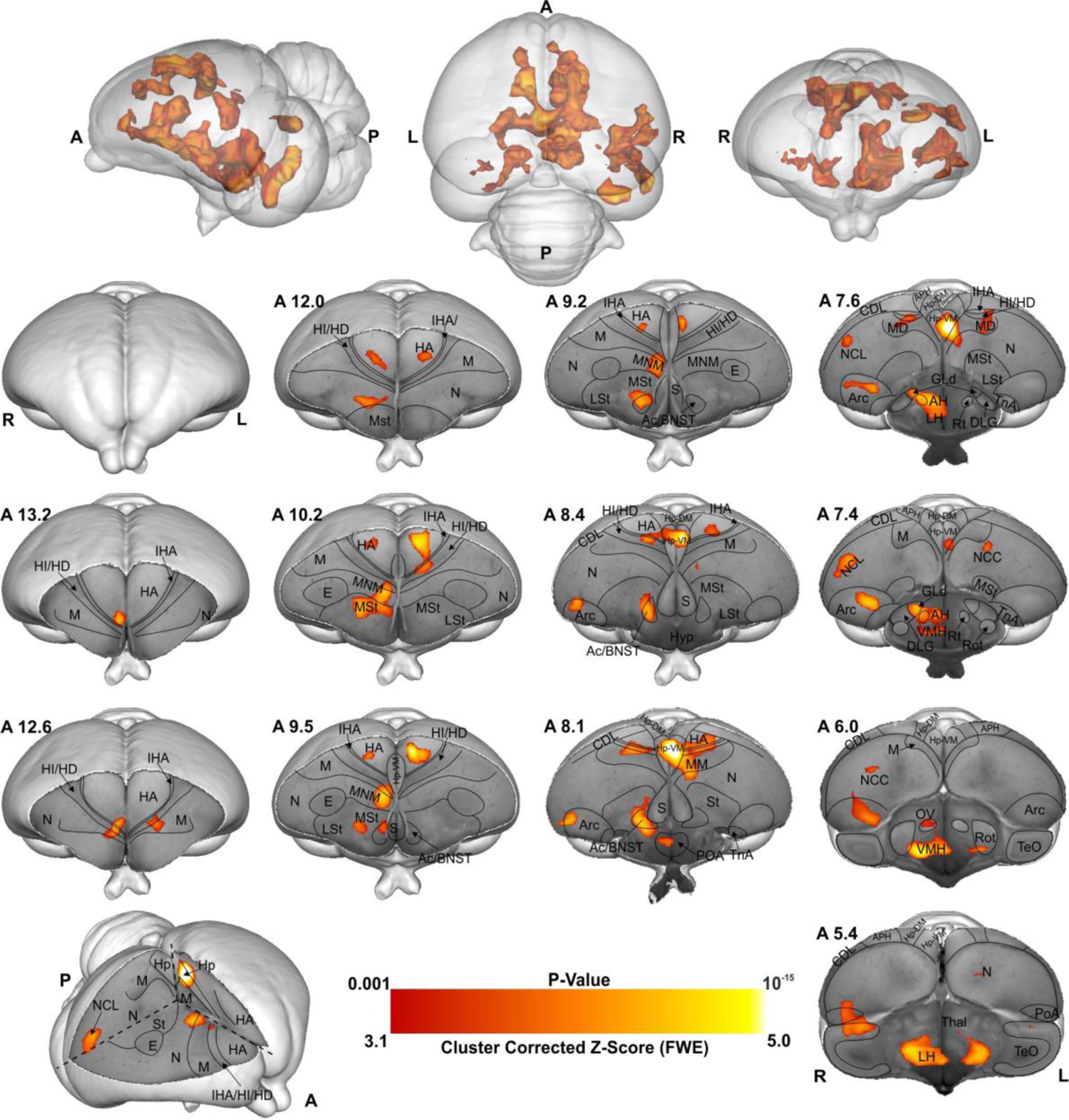
BOLD response pattern during imprinting memory retrieval. The high-resolution coronal slices at the different levels of an ex-vivo chick brain are in greyscale, while the contrast map represents the activation pattern during the presentation of the preferred imprinting object after imprinting learning has already occurred (Red group, the contrast of red light versus blue light conditions, n = 9, Z = 3.1 and p < 0.05 FEW corrected at the cluster level). The top row images show the 3D representation of the activation pattern inside a translucent chick brain. A 3D depiction of the chick brain is represented bottom left with an example window at the level of A 7.4. Anatomical borders (black lines) are based on the contrast difference of ex-vivo chick brain and Chick atlas (Herold et al., 2014; Kuenzel and Masson, 1988). The corresponding abbreviations of ROIs are listed in the Table S1.

Within the two interconnected *Social Behavior Network* and *Mesolimbic Reward System*, we detected significant bilateral activation in the ventromedial part of the hippocampus (Hp-VM), while rightward activation clusters in the bed nucleus of the stria terminalis (BNST), in the nucleus accumbens (Ac), in the medial striatum (MSt), in the medial and dorsal arcopallium (respectively AM and AD), in the posterior pallial amygdala (PoA) and in the preoptic, anterior and ventromedial areas of the hypothalamus (respectively POA, AH, and VMH).

## Discussion

Imprinting, a well-known form of early learning, has been widely used in the 70’s as a model to study the neurobiology of memory formation (reviews in Horn, 2004; Rose, 2000). Evidence for a crucial role played by the intermediate medial mesopallium (IMM) and NMm (jointly labelled as MNM) during the acquisition of imprinting memory was obtained. Further studies showed that the store in the IMM is only temporary, and that a transfer of information to another, unknown brain region, dubbed S’ (Honey et al., 1995), occurs after approximately 6 hours. These studies were conducted with either autoradiographic or lesion techniques and were unable to discover the full network of imprinting (Horn et al., 1979). Here we established a new fMRI protocol to study awake brain activity in newly hatched domestic chicks in order to discover the neural pathways of imprinting and the location of S’. After two days of imprinting training, with either a preferred (red) or a non-preferred (blue) colour (Maekawa et al., 2006), chicks were exposed to a sequence of the two stimulus colours inside the scanner. We provided evidence for a network of brain regions involved in the acquisition of a secondary imprinting memory (blue stimulus), and, in parallel, a neural network involved in the long-term storage of imprinting memory (red stimulus).

Visual information reaches the pallium both via the tecto- and the thalamofugal visual pathways. We observed a partial involvement of the nucleus rotundus (Rot), the thalamic link of the tectofugal pathway during acquisition (Fig. 5A). A rotundal involvement had already been reported in imprinted chicks (Harvey et al., 1998) and together with the present results, it suggests a minor tectofugal role during the early stages of imprinting learning. In contrast, the thalamofugal visual system seems to play a crucial role in processing imprinting information (as also reviewed in Nakamori et al., 2013). This pathway consists of the retinorecipient GLd (Aoki et al., 2015) that projects to the the interstitial nucleus of the hyperpallium apicale (IHA) of the visual Wulst, from where secondary projections reach the three pseudo-layers of the wulst hyperpallium densocellulare (HD), hyperpallium intercalatum (HI), and hyperpallium apicale (HA) (Stacho et al., 2020). We discovered both during memory formation and retrieval (Fig. 5B) significant activity patterns of all these thalamofugal components. Indeed, HD of dark-reared chicks exhibits topographically organised responses for red and blue objects (Maekawa et al., 2006). After imprinting on either one of the two colours, such organisation changes along the rostro-caudal axis showing imprinting-related plasticity already in the Wulst.

**Fig.5.**
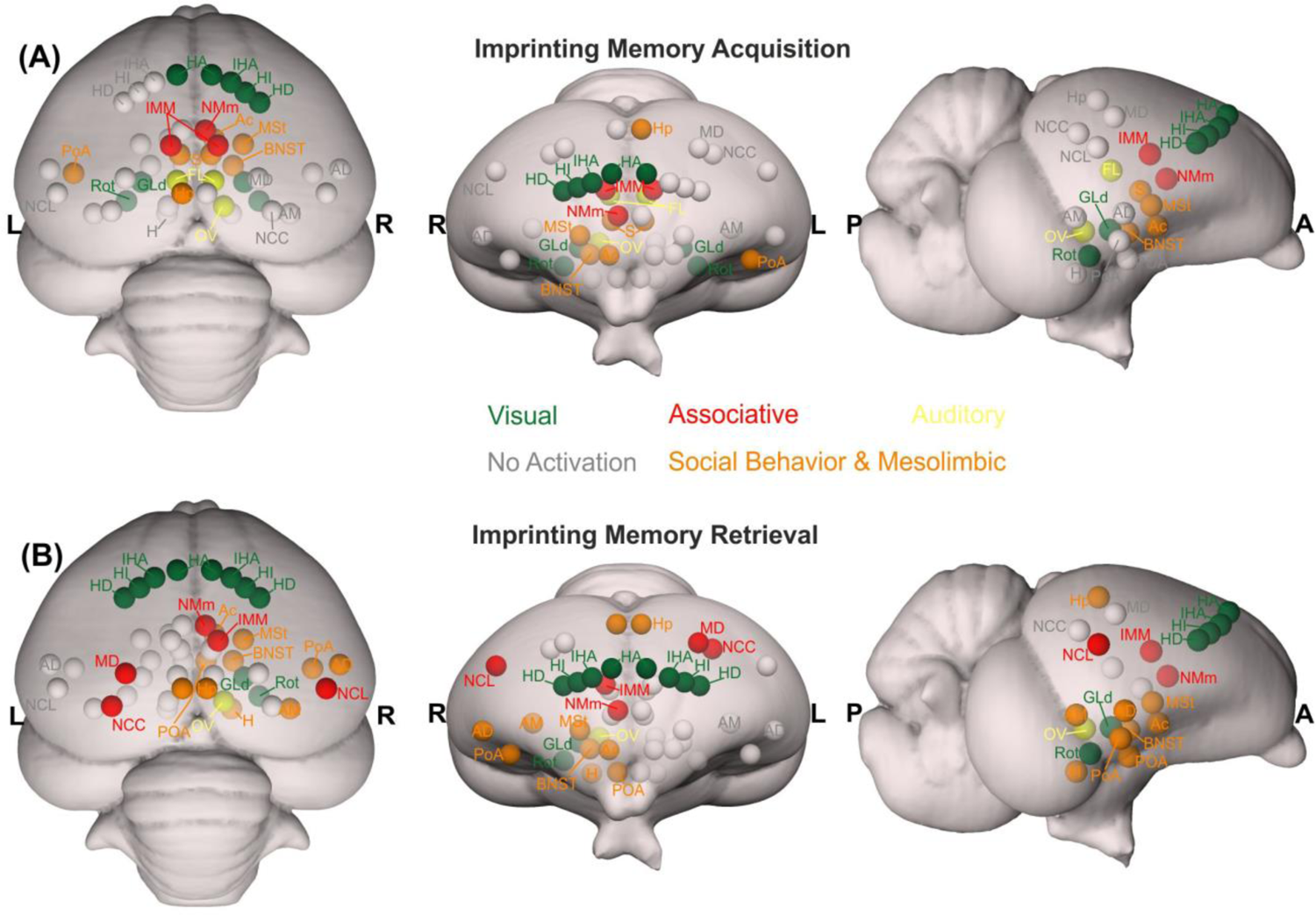
Schematic depiction of the activated prosencephalic areas during different phases of imprinting memory. (A) Network activated during imprinting memory acquisition are represented in colourful circles. (B) Network activated during imprinting memory retrieval are represented in colourful circles. The grey circles represent no activation. The corresponding abbreviations of ROIs are listed in the Table S1.

Previous studies showed that Wulst lesions lead to anterograde amnesia of visual imprinting memory (Maekawa et al., 2006). This possibly results from the loss of visual projections from HD to IMM (Nakamori et al., 2010; Shimizu et al., 1995), the associative medial pallial area that is crucial for the acquisition of imprinting memory (Horn, 2004). IMM projects back to HA, establishing a loop (Aoki et al., 2015). IMM, the ventrally located NMm and the nidopallium caudolaterale (NCL) have been shown to be involved during visual as well as auditory filial imprinting (Bock et al., 1997; Bredenkötter and Braun, 1997). Here we report a significant brain activation in IMM, NMm, and NCL during memory retrieval and, to a much lesser extent, in IMM and NMm during memory formation. Indeed, NMm and NCL undergo long-lasting synaptic changes after multimodal (visuo-auditory) imprinting training (Bock et al., 1997; Horn, 2004; Metzger et al., 1998). Imprinting training also impacts cell proliferation in NMm and NCL, but not in IMM (Komissarova and Anokhin, 2008). Thus, these three areas play important but differential roles in multimodal filial learning and the subsequent formation of long-term memory. Note that in the present study chicks were also exposed to the noise produced by the scanner. Thus, NMm and NCL on the one and the auditory n. ovoidalis (OV) – Field-L pathway on the other side, conceivably constituted the neural basis for the acoustic component of acquiring (blue group) or retrieving imprinting memory (red group).

But the interconnected higher associative regions, NMm and NCL do not only play a role for long-term memory-related mechanisms (Atoji and Wild, 2012; Behroozi et al., 2020; Metzger et al., 1998; Rook et al., 2021). NMm is also involved in sensorimotor learning and sequential behaviour (Helduser and Güntürkün, 2012), while NCL, largely accepted as a prefrontal-like field (von Eugen et al., 2020), is involved in working memory (Diekamp et al., 2002; Veit et al., 2014), executive control (Rose and Colombo, 2005) and in merging multi-sensory information in long-term memory engrams (Moll and Nieder, 2015). This evidence together with the present findings further supports the involvement of these regions in the long-term storage and flexible retrieval of a multimodal imprinting memory trace.

The motor output component of NMm and NCL is established by their projections to arcopallium and medial striatum (MSt, (Atoji and Wild, 2012; Csillag, 1999; Kröner and Güntürkün, 1999; Metzger et al., 2002; Schnabel et al., 1997)). Possibly, the initially pallially processed imprinting trace is thereby transferred into a striatum-dependent response strategy. As a result, striatal S-R associations are formed and once acquired, dominate the behaviour of the animal (Csillag, 1999). This also has been shown for passive avoidance learning. Here, the mnemonic nature of MSt (previously lobus paraolfactorius (Reiner et al., 2004)) goes hand in hand with that of IMM (Gibbs, 2008; Rose, 2000; Sojka et al., 1995), with increased density of synapses and dendritic spines being detectable some days after training in MSt, but not in IMM (Bradley et al., 1985; Sojka et al., 1995; Stewart and Rusakov, 1995). Additionally, after imprinting training glutamate receptor binding affinity increases both in MSt and arcopallium (Izawa et al., 2001; Johnston et al., 1993; McCabe and Horn, 1994), while, pre-imprinting arcopallial lesions impair memory acquisition (Lowndes et al., 1994).

We found enhanced brain activity in the most medial part of MSt both during acquisition and retrieval of imprinting memory, while for the dorsal and medial portions of arcopallium this was only observed for retrieval. These portions of MSt and arcopallium are enriched in the limbic system-associated membrane protein (LAMP, (Yamamoto and Reiner, 2005)). We also found a strong mesolimbic involvement in imprinting memory in the two interconnected *Social Behavior Network* and *Mesolimbic Reward System* (Goodson and Kingsbury, 2013; Newman, 1999; O’Connell and Hofmann, 2011). Here septum was involved only during memory formation. Arcopallium, preoptic area, anterior and ventromedial hypothalamus (POA, AH, VMH) were involved only during memory retrieval. In contrast, Hp, MSt, bed n. of the stria terminalis (BNST), n. accumbens (Ac) and posterior pallial amygdala (PoA) were involved during both memory formation and retrieval. While involvement of these systems in social predispositions associated with imprinting had already been observed (Lorenzi et al., 2017; Lorenzi and Vallortigara, 2021; Mayer et al., 2017), this is the first evidence for their involvement during imprinting. Such involvement could represent the motivational component linked to the association. Indeed, in the context of filial imprinting, emotional-motivational engagement must be particularly pronounced at different stages of the learning process. The septum seems to be preferentially involved during the first stages of imprinting and probably driving the chick’s attention toward salient predisposed moving stimuli. Previous studies also revealed septal involvement during the first exposure to a red object moving with abrupt changes of speed or an alive conspecific (Lorenzi et al., 2017; Mayer et al., 2017). Although BNST, Ac, MSt, and PoA seem to participate in both imprinting memory formation and retrieval, we found greater activity in the red group. Such enhanced activity may suggest a stronger emotional-motivational component after memory consolidation of the imprinting engram.

The HD of the Wulst has bidirectional connections with PoA and Hp (Atoji et al., 2018, 2006). We found a hippocampal (Hp) involvement both during imprinting memory formation and retrieval. The hippocampal formation is known for its role in memory in birds and mammals (Colombo and Broadbent, 2000). However, *c-fos* immunoreactivity in chicks revealed also a social role of Hp. The dorso- and ventromedial portions are involved in individual recognition in chicks (Corrales Parada et al., 2021). The same portions here were found to be involved in imprinting memory, strengthening a regional specialisation of hippocampus dedicated to social memory functions. Indeed, Hp projects ipsi- and contralaterally to IMM (Bradley et al., 1985) and is involved in filial imprinting (McCabe and Horn, 1994). We found a left Hp involvement during filial imprinting memory formation (blue group), and a bilateral one during memory retrieval (red group).

Interestingly, the brain activity pattern was predominantly right lateralised. Among the exceptions was a left Hp involvement during imprinting memory formation (blue group), and a bilateral Hp involvement during memory retrieval (red group). Lateralisation is a common feature in the avian brain, especially at different stages of memory formation (Moorman and Nicol, 2015; Rogers et al., 2013; Tulving et al., 1994). Right lateralisation during memory formation has been reported for passive avoidance learning (Lössner and Rose, 1983). Instead, for imprinting learning, time-shifts have been observed in the lateralisation pattern of IMM. The left IMM is involved at first in learning the features of the imprinting object, while the right IMM dominates during memory consolidation and the subsequent establishment of the long-term storage S’ (Cipolla-Neto et al., 1982; McCabe, 1991). A similar pattern of lateralisation has been proposed in the hemispheric encoding/retrieval asymmetry model (HERA) in humans, where the left hemisphere plays a dominant role during memory encoding and the right during retrieval (Tulving et al., 1994). Such evidence together leads to the hypothesis of a dual memory system for imprinting, in which different processes - acquisition and consolidation - take place in different hemispheres, with prominent right lateralisation for consolidation processes (Cipolla-Neto et al., 1982). Indeed, during memory consolidation, a glutamate injection into the right IMM disrupts imprinting memory, but it does not when injected into the left hemisphere (Johnston and Rogers, 1998). Our results may add a novel view on the idea of the dual memory system: While the visual thalamofugal nucleus GLd was bilaterally activated during acquisition, only the right side was active during retrieval. It is conceivable that right hemispheric memory consolidation increased top-down projections onto right sided sensory thalamic nuclei in order to focus attention on learned object properties (Kastner and Ungerleider, 2000). This then could activate and synchronize right hemispheric pallial areas according to attentional allocation, thereby inducing a right hemispheric superiority in imprinting memory retrieval (Saalmann et al., 2012).

Our findings could for the first time uncover a prosencephalic neural network that, among others, involves the *Social Behavior Network,* the *Mesolimbic Reward System,* and the medial meso-/nidopallium for long-term storage and retrieval of filial imprinting memory. As to be expected, the networks involved in memory formation and retrieval partially overlapped. However, network activity was more pronounced and further involved arcopallium and NCL in the retrieval condition. Thus, consolidation of imprinting memory results in a strengthening and expansion of the neural system that holds the engram in distributed manner. Within this perspective, the long searched site for imprinting memory dubbed as S’ by Gabriel Horn (Horn, 1985) is possibly this whole network within which the “prefrontal” NCL could be a central hub.

## Materials and methods

### Subjects

All procedures here presented followed all the applicable European Union and Italian laws, and guidelines for animals’ care and use and were approved by the Ethical Committee of the University of Trento OPBA and by the Italian Health Ministry (permit number 738/2019).

A local commercial hatchery (Azienda Agricola Crescenti, Brescia, Italy) provided fertilised eggs of the Aviagen Ross 308 strain (*Gallus gallus domesticus*). Eggs were incubated and hatched in the laboratory under controlled temperature (37.7°C) and humidity (60%) in darkness using FIEM MG140/200 Rural LCD EVO incubators. Soon after hatching, chicks were sexed by feather dimorphism, with a black cap on the head in order to prevent any visual stimulation. Twenty-six females were used in the present study. Females were used because they are known to exhibit stronger filial attachment with the imprinting object (Cailotto et al., 1989; Vallortigara, 1992; Vallortigara et al., 1990). Each chick underwent the experimental procedure only once. At the end of the experimental procedure, on post-hatching day 3, chicks were caged in groups with water and food *ad libitum,* at constant temperature (32.3°C) and with a 12:12 day-night light cycle until they were donated to local farmers.

### Imprinting

On the day of hatching, chicks were caged individually at a constant temperature of 32.3°C with water and food. In each cage (28×40×32 cm) the imprinting stimulus, a hollow plastic ball (diameter 3.5 cm), was suspended in the middle (7 cm from the floor, Fig. 1 A). Two optical fibres (diameter of 2mm) inserted in the ball were flickering at 5 Hz. Chicks prefer to imprint on a flickering than on a stationary light (James, 1959). For one group of chicks, the ball was flickering with red light (N = 9, dominant wavelength = 642 nm, intensity = 16.45 cd/m^2^,), for the other group with blue light (N = 8, dominant wavelength = 465 nm, intensity = 16.45 cd/m^2^). Being the only light provided in the environment, the established setup by Behroozi et al. (Behroozi et al., 2020) and a custom-written MATLAB code were used to automatically switch on and off the light, following a day-night cycle 12:12. During the daytime, to habituate the subjects to the noise of the scanner, a recording of the sound was provided twice per day, for a total amount of 5 hours per day, by two loudspeakers (Logitech) placed outside the cages.

### Acquisition and Pre-processing of fMRI data

All MRI experiments were recorded using a horizontal-bore small animal MRI scanner (7.0 T Bruker BioSpin, Ettlingen, Germany) equipped with a BGA-9 gradient set (380 mT/m, max. linear slew rate 3,420 T/m/s). A 72 mm transmit birdcage resonator was used for radio-frequency transmission. To reduce the motion artifacts resulting from body parts’ movements, a single-loop 20 mm surface coil was placed around the chicks’ head for signal reception.

### Localiser

At the beginning of each scanning session, a set of scout images (coronal, horizontal, and sagittal scans) were recorded as localisers to identify the position and orientation of the chick’s brain inside the MRI machine. The scout images were acquired using a multi-slice rapid acquisition (RARE) sequence with the following parameters: repetition time (TR) = 3000 ms, effective echo time (TE_eff_) = 41.2 ms, RARE factor = 32, N_average = 2, acquisition matrix = 128 × 128, the field of view (FoV) = 20 × 20 mm, spatial resolution = 0.156 × 0.156 mm^2^, slice thickness = 1 mm, number of slices = 8, slice orientation = coronal/horizontal/sagittal, with a total scan time of 18 s. This information has been used to position 9 coronal slices in a way (∼40° regarding coronal direction) to cover the entire telencephalon to record the fMRI time series.

### fMRI (task)

The blood-oxygen-level-dependent (BOLD) time series were recorded using a single-shot multi-slice RARE sequence adopted from Behroozi et al. (Behroozi et al., 2020, 2018) with the following parameters: TR/TE_eff_ = 4000/51.04 ms, RARE factor = 42, acquisition matrix = 64 × 64, FoV = 30 × 30 mm^2^, 9 coronal slices no gap between slices, slice thickness = 1 mm, slice order = interleaved. Since the eyes’ size is comparable to brain’s one, two saturation slices were manually positioned on the eyes to saturate the possible eye movement artifacts, which can corrupt the BOLD signal. A total of 540 volumes were recorded for each animal.

### fMRI (Rest)

Whole-brain resting-state fMRI data (200 volumes) of nine chicks were recorded using a single-shot RARE sequence with the same parameter as the task fMRI sequence.

### Structural MRI

High-resolution anatomical images were acquired using a RARE sequence with following parameters: TR/TE_eff_ = 6000/42.04 ms, RARE factor = 16, N_Average = 4, acquisition matrix = 160 × 160, FoV = 30 × 30 mm^2^, 39 coronal slices with no gap between slices, slice thickness = 0.33 mm, total scan time = 4 min.

### Experimental Task

Inside the fMRI machine, chicks were presented with two different stimulus types, imprinted (red/blue) and control colour (blue/red) with the same wavelength and intensity as the training phase. The light stimuli were generated using the established setup by Behroozi et al. (Behroozi et al., 2020). Stimuli were presented in a pseudo-random order in an ON/OFF block design experiment. The duration of ON blocks was 16 s. ON blocks were interleaved with a rest period of 24 s (OFF blocks, inter-trial interval (ITI)). In total, 48 trials were recorded during an fMRI session from each animal (24 trials per stimulus).

### Apparatus

A critical issue during awake fMRI scanning of animals is motion artifacts. Therefore, immobilisation of the animal’s head is essential to acquire an accurate fMRI time series. To this end, awake chicks were immobilised in a nonmagnetic custom-made restrainer, composed of a beak holder, blocks of plasticine around the head to immobilise it in a comfortable way, and a round RF coil on top of the head (Fig. 1B). Before the head fixation, the animal’s body was wrapped in paper tissue to prevent the other body parts’ movement (such as wings and feet) to avoid any possible motion artifacts. The animal’s body inside the paper tissue was tapped to the main body of the restrainer using a piece of medical tape.

### fMRI data processing

All BOLD time series were pre-processed using the FMRIB Software Library (FSL, version 6.0.4, https://fsl.fmrib.ox.ac.uk/fsl/fslwiki), the Analysis of Functional NeuroImages (AFNI, version 20.0.09 https://afni.nimh.nih.gov/), and Advanced Normalization Tools (ANTs, http://stnava.github.io/ANTs/) software. We performed the following pre-processing steps for each run: (i) converting dicom files to nifti format (using dcm2niix function); (ii) upscaling the voxel size by a factor of 10 (using AFNI’s 3drefit); (iii) discarding the first 5 volumes to ensure longitudinal magnetization reached steady state; (iv) motion correction using MCFLIRT (which aligns each volume to the middle volume of each run); (v) slice time correction to account for the long whole-brain acquisition time (4000 ms); interleaved acquisitions); (vi) despiking using 3dDespike algorithm in AFNI; (vii) removing non-brain tissue (using BET and manual cleaning); (viii) spatial smoothing with FWHM = 8 mm (using FSL’s SUSAN, after upscaling voxel size by factor of 10); (ix) global intensity normalization with grand mean = 10000 across scanning sessions for group analysis; (x) high-pass temporal filtering to remove slow drifts (cut-off at 100s); (xi) anatomical brain extraction (using BET function and cleaned manually); (xii) registration of the functional data to the high-resolution structural images using affine linear registration (FLIRT function, six degrees of freedom). For spatial normalization, a population-based template was constructed using antsMultivariateTemplateConstruction.sh script (ANTs). FMRIB’s Nonlinear Image Registration Tool (FNRIT) (Andersson et al., 2007) was used to spatially normalize the single subject anatomical images to the population-based template as a standard space. The head motion of animals was quantified using framewise displacement (FD) (Power et al., 2014). Three animals’ data were excluded due to the excessive head motion (over 20% of volumes were contaminated with FD > 0.2mm). For the remaining animals, the detected motion outliers were modeled as confound regressors during the general linear model (GLM) analysis to reduce the impact of head motion.

### General linear model (GLM) analysis

Whole-brain statistical analysis was performed using the FEAT (FMRI Expert Analysis Tool) to assess stimulus-evoked activation patterns. Single-subject GLM analysis was carried out to convolve the established double-gamma avian hemodynamic response function in pigeon brain by Behroozi et al. (Behroozi et al., 2020) (the closest brain in the structural organization to the chick brain) to the explanatory variables (on/off stimulation). The GLM contained the following three explanatory variables (EVs) and their temporal derivatives: (i) imprinting trials (red/blue); (ii) control trials (blue/red); (iii) junk trials (the first 16 trials were used as habituation period to the real magnet environment). In addition, six estimated head motion parameters (three translations and three rotations) and outlier volumes detected based on the FD analysis were modelled as confound EVs to remove the residual motion artifacts. To perform group inference, subject-level parameter estimates were taken into the second-level analysis using the mixed-effect model (FLAME1+2) to produce group-level estimates of each condition. FLAME 1+2 cluster-based approach has been used to threshold the group-level statistical maps for contrasts of interest with a cluster defining voxel threshold of *p* < 0.001 (Z > 3.1) for red group and *p* < 0.01 (Z > 2.3) for blue group and Family Error Wise (FEW) cluster significance threshold of *p* = 0.05.

### Visualization

To visualize the results, we took advantage of the high-resolution anatomical image acquired for another study. Briefly, five post-mortem chick brains were scanned using a fast-low angle shot (FLASH) sequence with following parameters: TR/TE = 50/4 ms, N_average = 6, acquisition matrix = 400 × 400 × 500, voxels size = 0.05× 0.05 × 0.05 mm^3^, total scan time = 19 h 48 min. The population-based template was co-registered nonlinearly (using FNIRT) to the high-resolution anatomical image of the chick brain. The contrasts of interest, eventually, were non-linearly warped to the high-resolution anatomical image. MANGO software (http://ric.uthscsa.edu/mango/mango.html, version 4.1) was used for 3D visualization of the activation patterns. Surf Ice software (https://www.nitrc.org/projects/surfice/, version v1.0.20201102 64bit x86-64 Windows) was used for surface rendering the chick brains with overlays to illustrate activated networks during imprinting acquisition and retrieval memory.

## Author Contributions

**Conceptualization:** Onur Güntürkün and Giorgio Vallortigara.

**Experiment design:** Mehdi Behroozi, Elena Lorenzi, Onur Güntürkün and Giorgio Vallortigara.

**Data Collection:** Mehdi Behroozi, Elena Lorenzi, and Sepideh Tabrik

**Methodology:** Mehdi Behroozi and Sepideh Tabrik

**Resources:** Alessandro Gozzi and Giorgio Vallortigara

**Writing:** All authors

**Visualization:** Mehdi Behroozi, Elena Lorenzi, Onur Güntürkün, and Sepideh Tabrik.

## Data and material availability

fMRI data for the chick imprinting and resting-state fMRI are available at (will be published after publishing manuscript). FSL software (https://fsl.fmrib.ox.ac.uk/fsl/fslwiki/, version 6.0.4) and MATLAB (2020b, MathWorks, USA) were used to process fMRI and behavioural data, respectively. Related processing codes can be found at https://github.com/mehdibehroozi/Imprinting-fMRI. All data needed to evaluate the conclusions in the paper are present in the paper and/or the Supplementary Materials.

## Supporting information

**Figure S1.**
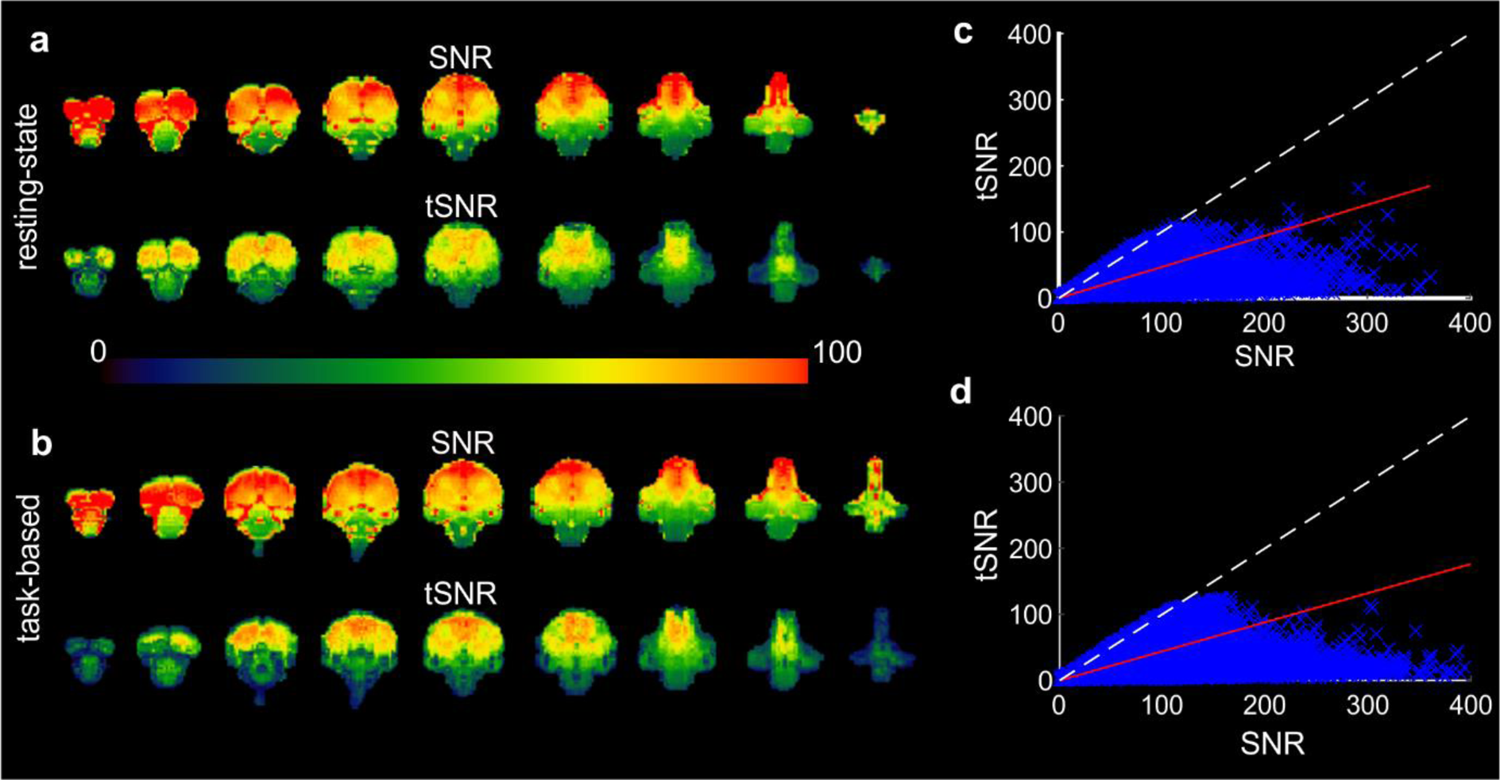
Temporal SNR and SNR assessments. (a,b) tSNR and SNR maps of resting-state and task-based fMRI data. 10 min resting-state data and an imprinting functional scan were used to calculate tSNR. The voxel-wise temporal SNR was defined as the voxel’s temporal mean divided by its temporal standard deviation. This figure illustrates that the tSNR (>50) is high in the regions of interest. (c,d) Overall SNR and tSNR were highly correlated with each other during resting-state and task-based fMRI data.

**Table S1.** Alphabetised list of abbreviations

| **Abb.** | **ROI name** | **Abb.** | **ROI name** |
| --- | --- | --- | --- |
| **Ac** | n. Accumbens | **LH** | Lateral Hypothalamus |
| **AD** | Dorsal Arcopallium | **LSt** | Lateral Striatum |
| **AH** | Anterior Hypothalamus | **M** | Mesopallium |
| **AM** | Medial Arcopallium | **MD** | Mesopallium Dorsale |
| **APH** | Area parahippocampalis | **MM** | Medial Mesopallium |
| **Arc** | Arcopallium | **NMm** | nidopallium medial pars medialis |
| **BNST** | Bed nucleus of the stria terminalis | **MNM** | IMM + MNm |
| **BOLD** | Blood-Oxygen-Level-Dependent | **MSt** | Medial Striatum |
| **CDL** | Area Corticoidea dorsolateralis | **N** | Nidopallium |
| **DLG** | Dorsolateral Geniculate n. | **NCL** | Nidopallium Caudolaterale |
| **FL** | Field-L | **NCC** | Central Caudal Nidopallium |
| **fMRI** | functional Magnetic Resonance | **OV** | n. ovoidalis |
| **E** | Entopallium | **POA** | Preoptic Area |
| **GLd** | n. Geniculatus lateralis pars dorsalis | **PoA** | n. posterior pallial amygdala |
| **HA** | Hyperpallium Apicale | **RARE** | rapid acquisition with relaxation |
| **HD** | Hyperpallium Densocellulare | **Rot** | n. Rotundus |
| **HI** | Hyperpallium Intercalatum | **Rt** | Reticular |
| **Hp** | Hippocampus | **S** | Septum |
| **Hp-** | Dorsomedial n. of the Hippocampus | **St** | Striatum |
| **Hp-** | Ventromedial n. of Hippocampus | **TeO** | Optic Tectum |
| **Hyp/H** | Hypothalamus | **Thal** | Thalamus |
| **IHA** | Interstitial n. of Hyperpallium Apicale | **TnA** | n. Taeniae Amygdalae |
| **IMM** | Intermediate Medial Mesopallium | **VMH** | Ventromedial hypothalamus |
| **ITI** | Inter trial interval |  |  |

## References

1. Andersson JLR, Jenkinson M, Smith SM. 2007. Non-linear optimisation. FMRIB technical report TR07JA1. Pr 16.

2. Aoki N, Yamaguchi S, Kitajima T, Takehara A, Katagiri-Nakagawa S, Matsui R, Watanabe D, Matsushima T, Homma KJ. 2015. Critical role of the neural pathway from the intermediate medial mesopallium to the intermediate hyperpallium apicale in filial imprinting of domestic chicks (Gallus gallus domesticus). Neuroscience 308:115–124. doi:10.1016/j.neuroscience.2015.09.014

3. Atoji Y, Saito S, Wild JM. 2006. Fiber connections of the compact division of the posterior pallial amygdala and lateral part of the bed nucleus of the stria terminalis in the pigeon (Columba livia). J Comp Neurol 499:161–182. doi:10.1002/cne.21042

4. Atoji Y, Sarkar S, Wild JM. 2018. Differential projections of the densocellular and intermediate parts of the hyperpallium in the pigeon (Columba livia). J Comp Neurol 526:146–165. doi:10.1002/cne.24328

5. Atoji Y, Wild JM. 2012. Afferent and efferent projections of the mesopallium in the pigeon (Columba livia). J Comp Neurol 520:717–741. doi:10.1002/cne.22763

6. Atoji Y, Wild JM. 2009. Afferent and efferent projections of the central caudal nidopallium in the pigeon (Columba livia). J Comp Neurol 517:350–370. doi:10.1002/cne.22146

7. Behroozi M, Chwiesko C, Ströckens F, Sauvage M, Helluy X, Peterburs J, Güntürkün O. 2018. In vivo measurement of T1 and T2 relaxation times in awake pigeon and rat brains at 7T. Magn Reson Med 79:1090–1100. doi:10.1002/mrm.26722

8. Behroozi M, Helluy X, Ströckens F, Gao M, Pusch R, Tabrik S, Tegenthoff M, Otto T, Axmacher N, Kumsta R, Moser D, Genc E, Güntürkün O. 2020. Event-related functional MRI of awake behaving pigeons at 7T. Nat Commun 11:1–12. doi:10.1038/s41467-020-18437-1

9. Behroozi M, Ströckens F, Helluy X, Stacho M, Güntürkün O. 2017. Functional Connectivity Pattern of the Internal Hippocampal Network in Awake Pigeons: A Resting-State fMRI Study. Brain Behav Evol 90:62–72. doi:10.1159/000475591

10. Boakes R, Panter D. 1985. Secondary imprinting in the domestic chick blocked by previous exposure to a live hen. Anim Behav 33:353–365. doi:10.1016/S0003-3472(85)80059-2

11. Bock J, Schnabel R, Braun K. 1997. Role of the dorso - caudal neostriatum in filial imprinting of the domestic chick: A pharmacological and autoradiographical approach focused on the involvement of NMDA-receptors. Eur J Neurosci 9:1262– 1272. doi:10.1111/j.1460-9568.1997.tb01481.x

12. Bradley P, Davies DC, Horn G. 1985. Connections of the hyperstriatum ventrale of the domestic chick (Gallus domesticus). J Anat 140:577–589.

13. Bradley P, Horn G, Bateson P. 1981. Imprinting - An electron microscopic study of chick hyperstriatum ventrale. Exp Brain Res 41:115–120. doi:10.1007/BF00236600

14. Bredenkötter M, Braun K. 2000. Development of Neuronal Responsiveness in the Mediorostral Neostriatum/Hyperstriatum Ventrale during Auditory Filial Imprinting in Domestic Chicks. Neurobiol Learn Mem 73:114–126. doi:https://doi.org/10.1006/nlme.1999.3923

15. Bredenkötter M, Braun K. 1997. Changes of neuronal responsiveness in the mediorostral neo striatum/hyperstriatum after auditory filial imprinting in the domestic chick. Neuroscience 76:355–365. doi:10.1016/S0306-4522(96)00381-8

16. Cailotto M, Vallortigara G, Zanforlin M. 1989. Sex differences in the response to social stimuli in young chicks. Ethol Ecol Evol 1:323–327. doi:10.1080/08927014.1989.9525502

17. Cipolla-Neto J, Horn G, McCabe BJ. 1982. Hemispheric asymmetry and imprinting: The effect of sequential lesions to the hyperstriatum ventrale. Exp Brain Res 48:22–27. doi:10.1007/BF00239569

18. Colombo M, Broadbent N. 2000. Is the avian hippocampus a functional homologue of the mammalian hippocampus? Neurosci Biobehav Rev. doi:10.1016/S0149-7634(00)00016-6

19. Corrales Parada CD, Morandi-Raikova A, Rosa-Salva O, Mayer U. 2021. Neural basis of unfamiliar conspecific recognition in domestic chicks (Gallus Gallus domesticus). Behav Brain Res 397:112927. doi:10.1016/j.bbr.2020.112927

20. Csillag A. 1999. Striato-telencephalic and striato-tegmental circuits: Relevance to learning in domestic chicks. Behav Brain Res 98:227–236. doi:10.1016/S0166-4328(98)00088-6

21. Csillag A, Montagnese CM. 2005. Thalamotelencephalic organization in birdsBrain Research Bulletin. Brain Res Bull. pp. 303–310. doi:10.1016/j.brainresbull.2005.03.020

22. Davey J, McCabe B, Horn G. 1987. Mechanisms of information storage after imprinting in the domestic chick. Behav Brain Res 26:209–210. doi:10.1016/0166-4328(87)90180-x

23. Diekamp B, Kalt T, Güntürkün O. 2002. Working memory neurons in pigeons. J Neurosci 22:RC210–RC210. doi:10.1523/jneurosci.22-04-j0002.2002

24. Gibbs ME. 2008. Memory systems in the chick: Regional and temporal control by noradrenaline. Brain Res Bull. doi:10.1016/j.brainresbull.2008.02.021

25. Goodson JL, Kingsbury MA. 2013. What’s in a name? Considerations of homologies and nomenclature for vertebrate social behavior networks. Horm Behav. doi:10.1016/j.yhbeh.2013.05.006

26. Güntürkün O, Karten HJ. 1991. An immunocytochemical analysis of the lateral geniculate complex in the pigeon (Columba livia). J Comp Neurol 314:721–749. doi:10.1002/cne.903140407

27. Harvey RJ, McCabe BJ, Solomonia RO, Horn G, Darlison MG. 1998. Expression of the GABAA receptor γ4-subunit gene: Anatomical distribution of the corresponding mRNA in the domestic chick forebrain and the effect of imprinting training. Eur J Neurosci 10:3024–3028. doi:10.1111/j.1460-9568.1998.00354.x

28. Helduser S, Güntürkün O. 2012. Neural substrates for serial reaction time tasks in pigeons. Behav Brain Res 230:132–143. doi:10.1016/j.bbr.2012.02.013

29. Herold C, Bingman VP, Ströckens F, Letzner S, Sauvage M, Palomero-Gallagher N, Zilles K, Güntürkün O. 2014. Distribution of neurotransmitter receptors and zinc in the pigeon (Columba livia) hippocampal formation: A basis for further comparison with the mammalian hippocampus. J Comp Neurol 522:2553–2575. doi:10.1002/CNE.23549

30. Honey RC, Horn G, Bateson P, Walpole M. 1995. Functionally Distinct Memories for Imprinting Stimuli: Behavioral and Neural Dissociations. Behav Neurosci 109:689– 698. doi:10.1037/0735-7044.109.4.689

31. Horn G. 2004. Pathways of the past: The imprint of memory. Nat Rev Neurosci. doi:10.1038/nrn1324

32. Horn G. 1985. Memory, imprinting, and the brain: an inquiry into mechanisms 10:315.

33. Horn G, Bradley P, McCabe BJ. 1985. Changes in the structure of synapses associated with learning. J Neurosci 5:3161–3168. doi:10.1523/jneurosci.05-12-03161.1985

34. Horn G, McCabe BJ, Bateson PPG. 1979. An autoradiographic study of the chick brain after imprinting. Brain Res 168:361–373. doi:10.1016/0006-8993(79)90176-8

35. Horn G, McCabe BJ, Cipolla-Neto J. 1983. Imprinting in the domestic chick: The role of each side of the hyperstriatum ventrale in acquisition and retention. Exp Brain Res 53:91–98. doi:10.1007/BF00239401

36. Izawa EI, Yanagihara S, Atsumi T, Matsushima T. 2001. The role of basal ganglia in reinforcement learning and imprinting in domestic chicks. Neuroreport 12:1743– 1747. doi:10.1097/00001756-200106130-00045

37. James H. 1959. Flicker: An unconditioned stimulus for imprinting. Can J Psychol Can Psychol 13:59–67. doi:10.1037/h0083767

38. Jarvis E, Güntürkün O, Bruce L, Csillag A, Karten H, Kuenzel W, Medina L, Paxinos G, Perkel DJ, Shimizu T, Striedter G, Martin Wild J, Ball GF, Dugas-Ford J, Durand SE, Hough GE, Husband S, Kubikova L, Lee DW, Mello C V., Powers A, Siang C, Smulders T V., Wada K, White SA, Yamamoto K, Yu J, Reiner A, Butler AB. 2005. Avian brains and a new understanding of vertebrate brain evolution. Nat Rev Neurosci. doi:10.1038/nrn1606

39. Johnston AN, Rogers LJ, Johnston GAR. 1993. Glutamate and imprinting memory: the role of glutamate receptors in the encoding of imprinting memory. Behav Brain Res 54:137–143. doi:10.1016/0166-4328(93)90072-X

40. Johnston ANB, Rogers LJ. 1998. Right hemisphere involvement in imprinting memory revealed by glutamate treatment. Pharmacol Biochem Behav 60:863–871. doi:10.1016/S0091-3057(98)00073-2

41. Karten HJ. 1967. The organization of the ascending auditory pathway in the pigeon (Columba livia) I. Diencephalic projections of the inferior colliculus (nucleus mesencephali lateralis, pars dorsalis). Brain Res 6:409–427. doi:10.1016/0006-8993(67)90055-8

42. Kastner S, Ungerleider LG. 2000. Mechanisms of visual attention in the human cortex. Annu Rev Neurosci. doi:10.1146/annurev.neuro.23.1.315

43. Komissarova N V., Anokhin K V. 2008. Effects of an imprinting procedure on cell proliferation in the chick brain. Neurosci Behav Physiol 38:289–296. doi:10.1007/s11055-008-0041-z

44. Kröner S, Güntürkün O. 1999. Afferent and efferent connections of the caudolateral neostriatum in the pigeon (Columba uvia): A retro- and anterograde pathway tracing study. J Comp Neurol 407:228–260. doi:10.1002/(SICI)1096-9861(19990503)407:2<228::AID-CNE6>3.0.CO;2-2

45. Kuenzel WJ, Masson M. 1988. A stereotaxic atlas of the brain of the chick (Gallus domesticus). Johns Hopkins University Press.

46. Lemche E. 2020. Research evidence from studies on filial imprinting, attachment, and early life stress: a new route for scientific integration. Acta Ethol. doi:10.1007/s10211-020-00346-7

47. Lorenz K. 1935. Der Kumpan in der Umwelt des Vogels. J für Ornithol 1935 832 83:137–213. doi:10.1007/BF01905355

48. Lorenzi E, Mayer U, Rosa-Salva O, Vallortigara G. 2017. Dynamic features of animate motion activate septal and preoptic areas in visually naïve chicks (Gallus gallus). Neuroscience 354:54–68. doi:10.1016/j.neuroscience.2017.04.022

49. Lorenzi E, Vallortigara G. 2021. Evolutionary and neural bases of the sense of animacy In: Kaufman AB, Call J, Kaufman JC, editors. The Cambridge Handbook of Animal Cognition. Cambridge University Press.

50. Lössner B, Rose SPR. 1983. Passive Avoidance Training Increases Fucokinase Activity in Right Forebrain Base of Day-Old Chicks. J Neurochem 41:1357–1363. doi:10.1111/j.1471-4159.1983.tb00833.x

51. Lowndes M, Davies DC, Johnson MH. 1994. Archistriatal Lesions Impair the Acquisition of Filial Preferences During Imprinting in the Domestic Chick. Eur J Neurosci 6:1143–1148. doi:10.1111/j.1460-9568.1994.tb00612.x

52. Maekawa F, Komine O, Sato K, Kanamatsu T, Uchimura M, Tanaka K, Ohki-Hamazaki H. 2006. Imprinting modulates processing of visual information in the visual wulst of chicks. BMC Neurosci 2006 71 7: 1–13. doi:10.1186/1471-2202-7-75

53. Mayer U, Rosa-Salva O, Vallortigara G. 2017. First exposure to an alive conspecific activates septal and amygdaloid nuclei in visually-naïve domestic chicks (Gallus gallus). Behav Brain Res 317:71–81. doi:10.1016/j.bbr.2016.09.031

54. McCabe BJ. 2013. Imprinting. Wiley Interdiscip Rev Cogn Sci 4:375–390. doi:10.1002/wcs.1231

55. McCabe BJ. 1991. Hemispheric asymmetry of learning-induced changesNeural and Behavioural Plasticity. Oxford University Press. pp. 262–276. doi:10.1093/acprof:oso/9780198521846.003.0010

56. McCabe BJ, Horn G. 1994. Learning-related changes in Fos-like immunoreactivity in the chick forebrain after imprinting. Proc Natl Acad Sci U S A 91:11417–11421. doi:10.1073/pnas.91.24.11417

57. Metzger M, Jiang S, Braun K. 2002. A quantitative immuno-electron microscopic study of dopamine terminals in forebrain regions of the domestic chick involved in filial imprinting. Neuroscience 111:611–623. doi:10.1016/S0306-4522(01)00611-X

58. Metzger M, Jiang S, Braun K. 1998. Organization of the dorsocaudal neostriatal complex: A retrograde and anterograde tracing study in the domestic chick with special emphasis on pathways relevant to imprinting. J Comp Neurol 395:380–404. doi:10.1002/(SICI)1096-9861(19980808)395:3<380::AID-CNE8>3.0.CO;2-Z

59. Moll FW, Nieder A. 2015. Cross-modal associative mnemonic signals in crow endbrain neurons. Curr Biol 25:2196–2201. doi:10.1016/j.cub.2015.07.013

60. Moorman S, Nicol AU. 2015. Memory-related brain lateralisation in birds and humans. Neurosci Biobehav Rev. doi:10.1016/j.neubiorev.2014.07.006

61. Müller-Schwarze D, Müller-Schwarze C. 1971. Olfactory imprinting in a precocial mammal. Nature 229:55–56. doi:10.1038/229055a0

62. Nakamori T, Maekawa F, Sato K, Tanaka K, Ohki-Hamazaki H. 2013. Neural basis of imprinting behavior in chicks. Dev Growth Differ. doi:10.1111/dgd.12028

63. Nakamori T, Sato K, Atoji Y, Kanamatsu T, Tanaka K, Ohki-Hamazaki H. 2010. Demonstration of a neural circuit critical for imprinting behavior in chicks. J Neurosci 30:4467–4480. doi:10.1523/JNEUROSCI.3532-09.2010

64. Newman SW. 1999. The medial extended amygdala in male reproductive behavior. A node in the mammalian social behavior network. Ann N Y Acad Sci 877:242–257. doi:10.1111/j.1749-6632.1999.tb09271.x

65. O’Connell LA, Hofmann HA. 2011. The Vertebrate mesolimbic reward system and social behavior network: A comparative synthesis. J Comp Neurol. doi:10.1002/cne.22735

66. Power JD, Mitra A, Laumann TO, Snyder AZ, Schlaggar BL, Petersen SE. 2014. Methods to detect, characterize, and remove motion artifact in resting state fMRI. Neuroimage 84:320–341. doi:10.1016/j.neuroimage.2013.08.048

67. Reiner A, Perkel DJ, Bruce LL, Butler AB, Csillag A, Kuenzel W, Medina L, Paxinos G, Shimizu T, Striedter G, Wild M, Ball GF, Durand S, Gütürkün O, Lee DW, Mello C V., Powers A, White SA, Hough G, Kubikova L, Smulders T V., Wada K, Dugas-Ford J, Husband S, Yamamoto K, Yu J, Siang C, Jarvis ED. 2004. Revised Nomenclature for Avian Telencephalon and Some Related Brainstem Nuclei. J Comp Neurol. doi:10.1002/cne.20118

68. Rogers LJ, Vallortigara G, Andrew RJ. 2013. Divided brains: The biology and behaviour of brain asymmetries. Cambridge University Press. doi:10.1017/CBO9780511793899

69. Rook N, Tuff JM, Isparta S, Masseck OA, Herlitze S, Güntürkün O, Pusch R. 2021. AAV1 is the optimal viral vector for optogenetic experiments in pigeons (Columba livia). Commun Biol 4:1–16. doi:10.1038/s42003-020-01595-9

70. Rose J, Colombo M. 2005. Neural correlates of executive control in the avian brain. PLoS Biol 3:1139–1146. doi:10.1371/journal.pbio.0030190

71. Rose SPR. 2000. God’s organism? The chick as a model system for memory studies. Learn Mem. doi:10.1101/lm.7.1.1

72. Saalmann YB, Pinsk MA, Wang L, Li X, Kastner S. 2012. The pulvinar regulates information transmission between cortical areas based on attention demands. Science (80-) 337:753–756. doi:10.1126/science.1223082

73. Salzen EA, Lily RE, McKeown JR. 1971. Colour preference and imprinting in domestic chicks. Anim Behav 19:542–547. doi:10.1016/S0003-3472(71)80109-4

74. Schnabel R, Metzger M, Jiang S, Hemmings HC, Greengard P, Braun K. 1997. Localization of dopamine D1 receptors and dopaminoceptive neurons in the chick forebrain. J Comp Neurol 388:146–168. doi:10.1002/(SICI)1096-9861(19971110)388:1<146::AID-CNE10>3.0.CO;2-T

75. Shanahan M, Bingman VP, Shimizu T, Wild M, Güntürkün O. 2013. Large-scale network organisation in the avian forebrain: A connectivity matrix and theoretical analysis. Front Comput Neurosci 7. doi:10.3389/fncom.2013.00089

76. Shimizu T, Cox K, Karten HJ. 1995. Intratelencephalic projections of the visual wulst in pigeons (Columba livia). J Comp Neurol 359:551–572. doi:10.1002/cne.903590404

77. Sojka M, Davies HA, Harrison E, Stewart MG. 1995. Long-term increases in synaptic density in chick CNS after passive avoidance training are blocked by an inhibitor of protein synthesis. Brain Res 684:209–214. doi:10.1016/0006-8993(95)00403-D

78. Spalding DA. 1954. Instinct: With original observations on young animals. Br J Anim Behav 2:2–11. doi:10.1016/S0950-5601(54)80075-X

79. Stacho M, Herold C, Rook N, Wagner H, Axer M, Amunts K, Güntürkün O. 2020. A cortex-like canonical circuit in the avian forebrain. Science (80-) 369. doi:10.1126/science.abc5534

80. Stewart MG, Rusakov DA. 1995. Morphological changes associated with stages of memory formation in the chick following passive avoidance training. Behav Brain Res 66:21–28. doi:10.1016/0166-4328(94)00119-Z

81. Tulving E, Kapur S, Craik FIM, Moscovitch M, Houle S. 1994. Hemispheric encoding/retrieval asymmetry in episodic memory: Positron emission tomography findings. Proc Natl Acad Sci U S A 91:2016–2020. doi:10.1073/pnas.91.6.2016

82. Vallortigara G. 2021. Born Knowing: Imprinting and the Origins of Knowledge. The MIT Press. doi:10.7551/mitpress/14091.001.0001

83. Vallortigara G. 1992. Affiliation and aggression as related to gender in domestic chicks (Gallus gallus). J Comp Psychol 106:53–57. doi:10.1037/0735-7036.106.1.53

84. Vallortigara G, Cailotto M, Zanforlin M. 1990. Sex differences in social reinstatement motivation of the domestic chick (Gallus gallus) revealed by runway tests with social and nonsocial reinforcement. J Comp Psychol 104:361–367. doi:10.1037/0735-7036.104.4.361

85. Veit L, Hartmann K, Nieder A. 2014. Neuronal correlates of visual working memory in the corvid endbrain. J Neurosci 34:7778–7786. doi:10.1523/JNEUROSCI.0612-14.2014

86. Vicedo M. 2013. The Nature and Nurture of Love. University of Chicago Press. doi:10.7208/chicago/9780226020693.001.0001

87. von Eugen K, Tabrik S, Güntürkün O, Ströckens F. 2020. A comparative analysis of the dopaminergic innervation of the executive caudal nidopallium in pigeon, chicken, zebra finch, and carrion crow. J Comp Neurol cne. 24878. doi:10.1002/cne.24878

88. Yamamoto K, Reiner A. 2005. Distribution of the limbic system-associated membrane protein (LAMP) in pigeon forebrain and midbrain. J Comp Neurol 486:221–242. doi:10.1002/cne.20562

89. Yamamoto K, Sun Z, Hong BW, Reiner A. 2005. Subpallial amygdala and nucleus taeniae in birds resemble extended amygdala and medial amygdala in mammals in their expression of markers of regional identity. Brain Res Bull 66:341–347. doi:10.1016/j.brainresbull.2005.02.016

